# siland: an R package for estimating the spatial influence of landscape

**DOI:** 10.1101/692566

**Authors:** Florence Carpentier, Olivier Martin

## Abstract

**Context:** The spatial distributions of species and populations are both influenced by local variables and by characteristics of surrounding landscapes. Understanding how landscape features spatially structure the frequency of a trait in a population, the abundance of a species or the species’ richness remains difficult specially because the spatial scale effects of the landscape variables are often unknown.

**Objectives:** Here, we present “siland”, an R package for analyzing the effect of landscape features on georeferenced point observations (described in a Geographic Information System shapefile format).

**Methods & Results:** “siland” simultaneously estimates the spatial scales and intensities of landscape variable effects. It does not require any information about the scale of effect. Two methods are available: one is based on focal sample site (*Bsiland method*, b for buffer) and one is distance weighted using Spatial Influence Function (*Fsiland method*, f for function). ‘siland’ allows for effects tests, effects maps and models comparison.

**Conclusions:** Adaptable and user-friendly, the “siland” package is a very practical tool to perform landscape analysis.

## Introduction

Numerous studies demonstrate that the distribution of species richness and abundance depend on both local and landscape variables (García et al., 2011; Rusch et al., 2016; Remm et al., 2017). However, studying the relationships between landscape and species distributions remains challenging because the shape and the scale of landscape effects are unknown (Miguet et al., 2016) and can be missed if assessed at an incorrect scale (Smith et al., 2011). The studied data usually contains georeferenced observations at point sites, named response variables hereafter, and the description of several landscape spatial variables, named landscape variables hereafter. Their studies are often referred to as focal patch studies (Thornton et al., 2011). To identify the scale of landscape variable effects, the common approach consists of the following: (i) *a priori* defining a set of scales; (ii) creating summary variables by computing measures of the landscape variables within discs or rings of radii equal to each scale centered on the observation sites (named buffers hereafter); and (iii) applying a regression model to the response variable with the summary variables as the explanatory variables, for example, a linear model or a random forest algorithm (Bradter et al., 2012). For each landscape variable, the scale of effect is then considered to be the size of the buffer best explaining the response variable.

The main disadvantage of this method is that the number of explanatory variables artificially increases with the number of spatial scales considered. One then faces a complex statistical dilemma, which is dealing with numerous explanatory variables that by their construction are highly correlated. Consequently, the potential scales chosen are often too few and their ranges are too limited (Jackson & Fahrig, 2015). Finally, the effect of a landscape variable is modelled as uniform within the buffer and as null outside it (Chandler & Hepinstall-Cymerman, 2016), which is unrealistic and biologically unjustified as a continuously decreasing effect is expected (Moilanen & Hanski, 2001). Several new methods based on distance-weighted effects have been proposed to model a distance-decreasing effect (Aue et al., 2011; Henry et al., 2012; Serckx et al., 2016), but they explore a limited predefined set of spatial scales for predictors. Other methods quantified the scale of landscape effects without an *a priori* choice of spatial scales (Walsh & Webb, 2014; Chandler & Hepinstall-Cymerman, 2016). However, none of these methods are yet implemented in a ready-to-use software. Huais (2018) proposed a very convenient R function “multifit” to select scales but with some limitation (it is not distance weighted, requires the choice of a set of scales), while calling for further developments of such a method to generalise this type of automated analysis.

Here, we present “siland”, a package for the R statistical computing environment dedicated to landscape effect analysis. Two methods are available. Both estimate the scales of effect of each landscape variable using maximum likelihood estimation. In the first method, Bsiland method hereafter (B for buffer), the effect of a landscape variable is modeled as in classic methods based on focal sample site, i.e. considered as constant over a disc centered on the observation point. But contrarily to the previous method, it does not require a first definition of tested radii since the optimal buffer radii are estimated. In the second method the effect of landscape is based on a weighted distance, as in the framework proposed by Chandler & Hepinstall-Cymerman (2016). The decrease in weight with distance is modeled by a Spatial Influence Function (SIF). The parameter of the SIF defined the scale of effect of a landscape variable. In this second method, named Fsiland (F for function), the parameters of the SIFs are estimated for each landscape variable. The main functions of siland allow the user (i) to estimate the intensity of local and landscape effects and the scale parameter of each landscape variable, (ii) to test these effects and (iii) to plot landscape effects on maps. We exemplified the package use by analysing the landscape effects of conventional and organic orchards on the density of codling moth larvae per apple tree.

## Models and methods

We consider a response variable measured at *n* different sites denoted *Y_i_* (*i* stands for a site), *L* local variables which can be continuous or discrete and are denoted as 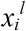 (*l* stands for a local variable and *i* for a site) and *K* landscape variables denoted as 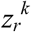 (*k* stands for a landscape variable and *r* for a polygon in the landscape). In the Bsiland method, the effect of landscape variables is modelled using buffers with 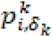, the percentage of the landscape variable *k* in a buffer of radius *δ_k_* and centered on site *i*. Since the Bsiland model is based on the generalized linear model framework of generalized linear models, the expected value of the response variable *Y_i_* is modelled as follows:

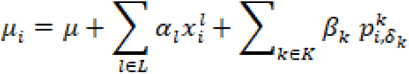

where *μ* is the intercept, α_1_ and β_k_ are the effects of local and landscape variables, respectively. All parameters, *μ, {*α_*1*,.._, α_*K*_*}*, *{*β_*1*.._, β_,*K*_*}* but also *{δ_1,.._, δ_K_}* radii of the buffers of the landscape variables are simultaneously estimated by likelihood maximization.

The Fsiland method is based on Spatial Influence Functions (SIFs) in a similar framework to Chandler & Hepinstall-Cymerman’s (2016). The entire study area is rasterized, *i.e*. pixelated on a regular grid, named *R*. The value of each landscape variable *k* at a pixel *r* is described in 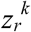. For instance, if the landscape variable *k* is a presence/absence variable, 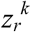 is equal to one or zero. The expected value of the response variable *Y_t_* is then modelled as follows:

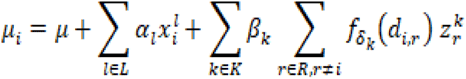

where *f_δk_(.)* is the **SIF** associated with the landscape variable *k* and *d_i,r_* is the distance between the center of pixel *r* and the observation at site *i*. The SIF is a density function decreasing with the distance. The scale of effect of a landscape variable *k* is calibrated through the parameter δ_*k*_, the mean distance of *f_δ_*. Two families of SIF are currently implemented in the siland package, exponential and Gaussian families defined as *f_δ_(d)*=*2/(πδ^2^)exp(−2d/δ)* and *f_δ_(d)*=*1/(2δ√π)exp(-dπ/2δ)^2^*, respectively (Austerlitz et al., 2004). The effect of a landscape variable *k* is modelled by two parameters: an intensity parameter, *β_k_* describing its strength and its direction and a scale parameter, *δ_k_*, describing how this effect declines with distance. Each pixel potentially has an effect on the response variable at any observation site. No set of scales of effects is initially determined.

## Package description

The siland package is written entirely in the scientific computing language R (R Core Team, 2019). It is available on CRAN (https://cran.r-project.org/web/packages/SILand/index.html) and new developments are available on https://github.com/silandpackage/siland. The analyses presented here were performed using the package siland 2.0.

## Case study

We illustrated the abilities of siland on an example previously described and analyzed in Ricci et al. (2009): the study of codling moth densities, an insect pest specialized on apple orchards in the Basse Durance Valley in southeastern France (see figure 1). The datasets can be extracted from the package using the commands `data(dataCmoth)` and `data(landCmoth)`. The complete analysis script and outputs are available in Supplementary Information (SI 1 and SI 2, respectively). `dataCmoth` is a data frame with two columns named `X` and `Y` containing the observation locations, a column `Cmoth`, containing the response variable of the study (the mean number of codling moths in the orchards), and a column `trait`, describing a local variable (the number of insecticide treatments applied in the orchard). `landCmoth` is a sf object (from package sf (Pebesma,2018)) describing the landscape variables: conventional tree orchards (`conv`), organic tree orchards (`org`) where `conv` is equal to 1 if the land use is associated to orchard with conventional practice and 0 otherwise, and so is it for `org`.

**Fig. 1:**
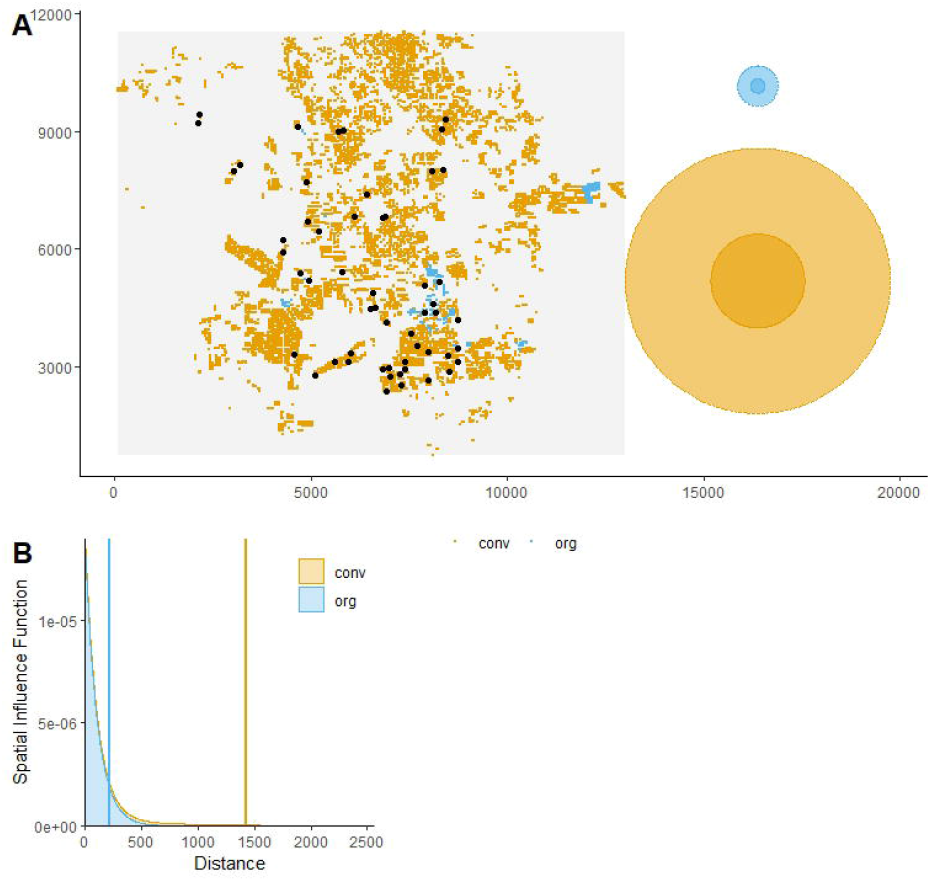
Map of the observations in the study site and the estimated spatial influence functions (SIFs) The figure A is obtained with `plotFsiland(resF,landCmoth,data=dataCmoth)`. Black points represent locations of response variable observations. Yellow and blue squares are the pixels where conventional and organic orchards respectively are present. At the right margin, the light and dark discs represent area of medium influence and significant influence, respectively. Blue and yellow discs represent conventional and the organic orchards, respectively. The figure B is obtained with `plotFsiland.sif(resF)`. The blue and orange lines represent the estimated SIF for the organic and conventional orchards, respectively. The vertical lines represent the SIF median.

## Main functions

The main functions are described in Table 1, while a full description of package functions is available at https://cran.r-project.org/web/packages/siland/siland.pdf. The package siland requires two objects containing data. The first object is a data frame composed by two columns named X and Y containing the observation locations, a third column representing the response variable and eventually other columns representing local variables. The second object is a sf object describing landscape variables. It can be obtained directly from shape files of landscape map by using the function `st_read()` of the package sf (see `vignette(siland)` for more details).

**Table 1.**
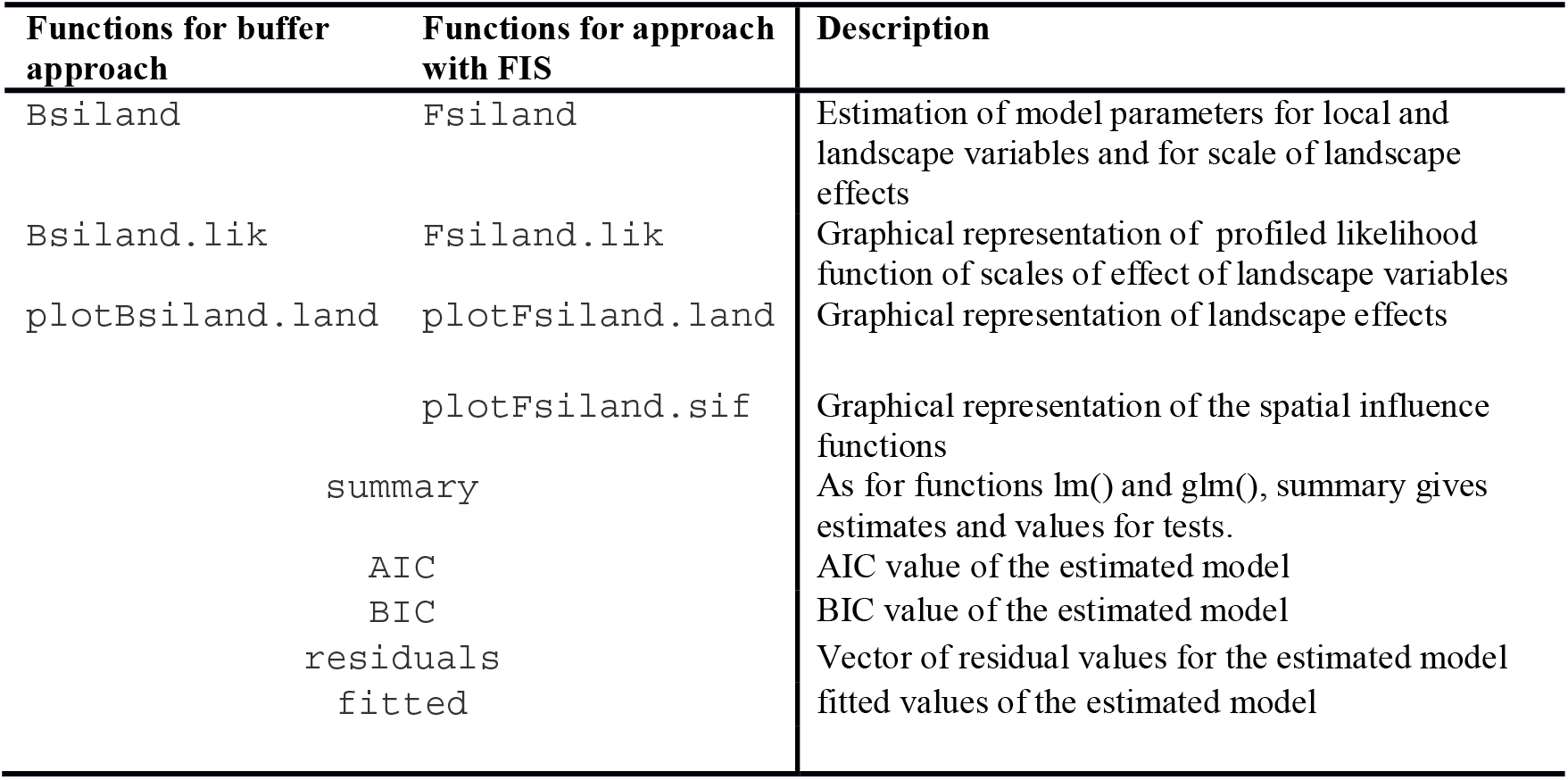
Main functions of siland package. Detailed information about these functions are given in https://cran.r-project.org/web/packages/siland/siland.pdf.

Model estimation is performed using the function `Bsiland()` for Bsiland method and the function `Fsiland()` for Fsiland method. The arguments of the both functions are similar: formula of the model, `land` the sf object describing landscape variables and `data`, the observation data frame. The syntax of the formula “y ~ model” is similar to `lm()` function of the stat package. In the model term of the formula, the explanatory variables are added using the symbol “+”. The explanatory variables described in the data frame data are considered as local in the model, those described in the sf object `land` are considered as landscape variables. Local effects can be modelled as fixed or random (using the syntax `(1|x)`, see `vignette (siland)` for more details). Interaction terms can be considered using the usual symbols “*” or “:”. Notice that only interactions between local x local and local x landscape variables are considered. Various types of response variables can be considered using the argument `family` which describes the assumed distribution of the response variable and can take the values `“gaussian”, “poisson”` and “`binomial`” for data of continuous, counting or presence-absence types, respectively (and their associated link functions identity, log and logit respectively).

Using the argument `border`, the spatial effect of landscape variable can be considered from the observation locations `(border=F)` or from the border of the polygon where observations are located `(border=T)` (see Figure 2). For `Fsiland()`, the additional argument `sif` indicates the family of SIF chosen `“exponential”` (by default) or `“gaussian”`.

**Fig. 2:**
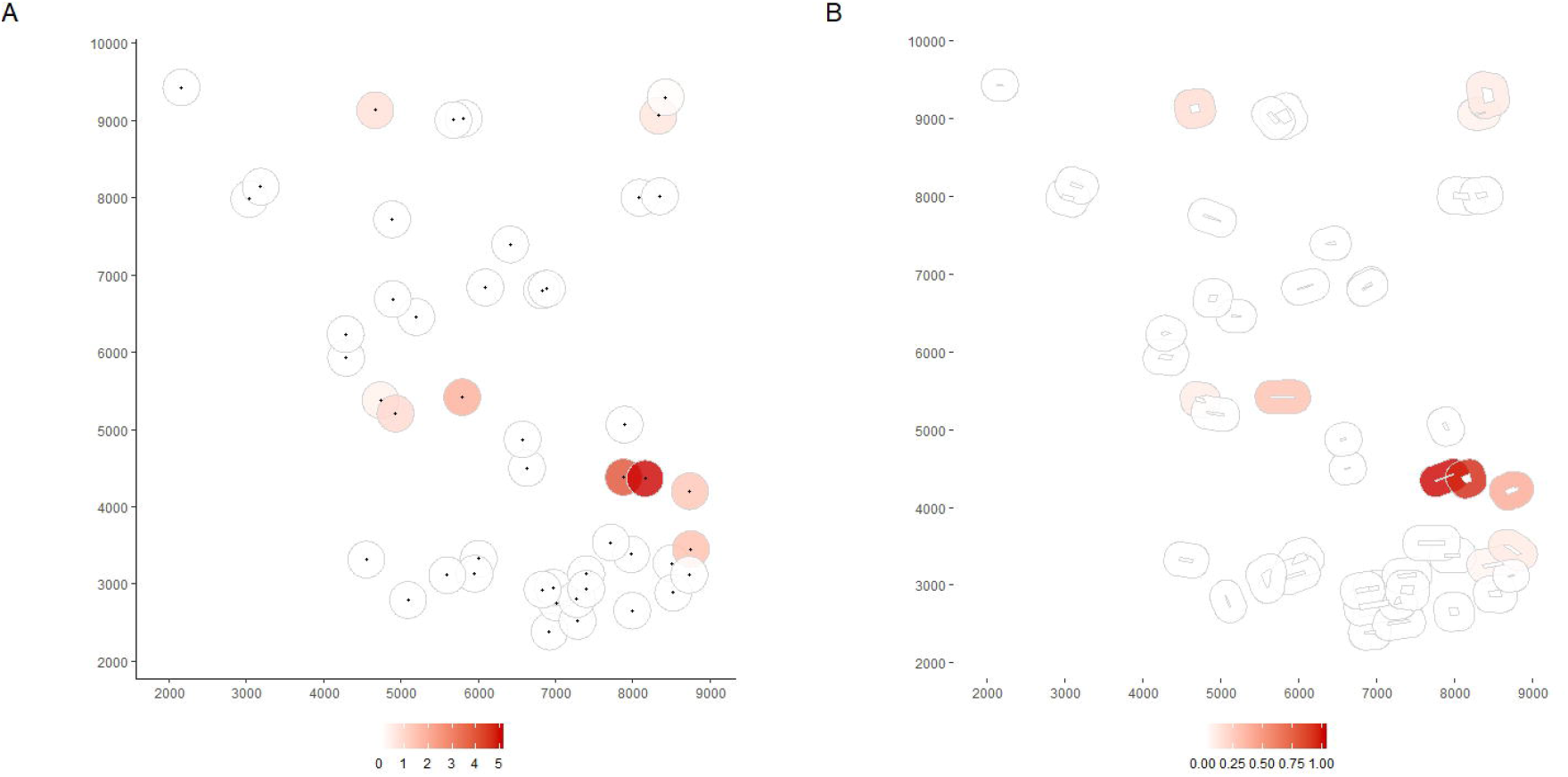
Maps of the predicted effect estimated by the Bsiland method obtained with the function plotBsiland.land(). The buffer model was estimated using the command: `resB2=Bsiland(Cmoth~trait+conv+org,land=landCmoth,data=dataCmoth,border=border)`. The colored area represents the buffer around each observation location estimated for the organic orchards effect. At the bottom margin, the bar color gives the effect intensity. Buffers are modeled in graphic A, from the observation locations `(border=FALSE)` and in graphic B from the border of the orchard of each observation `(border=TRUE)`.

The object returned by the functions `Bsiland()` and `Fsiland()` displays the parameters estimation and a test of the global landscape effect. The function `summary()` applied on the result object provides significance tests of the intensity of the effect of explanatory variables (local or landscape).

The two methods are based on likelihood maximization. The functions `Bsiland.lik()` and `Fsiland.lik()` aim to point out some optimization problems. They provide representations of the minus loglikelihood in function of buffer radius or mean distance of the SIF, respectively (see SI 1). Values of minus loglikelihood lower than the estimated one indicate that the estimation procedure did not perform correctly. In such cases, the estimation needs to be reiterated using different initial values of effects scales (with argument `init` of functions `Fsiland()` and `Bsiland()`).

## Graphics outputs

The package siland proposes various functions for graphic representation of estimated landscape effects. The function “`plotFsiland.land`” displays landscape effect map for the estimations of the Fsiland method (Figure 3). The influence of each landscape variable is plotted over all the study area. The global influence of the landscape, i.e. the sum of the effects of all landscape variables can also be plotted. The function `plotBsiland.land()` displays landscape effect map for the estimation of the Bsiland method (Figure 2). For each landscape variable, the estimated area of effect (buffer of estimated radius) is plotted around each observation locations. The color of the area represents the intensity of the effect.

**Fig. 3:**
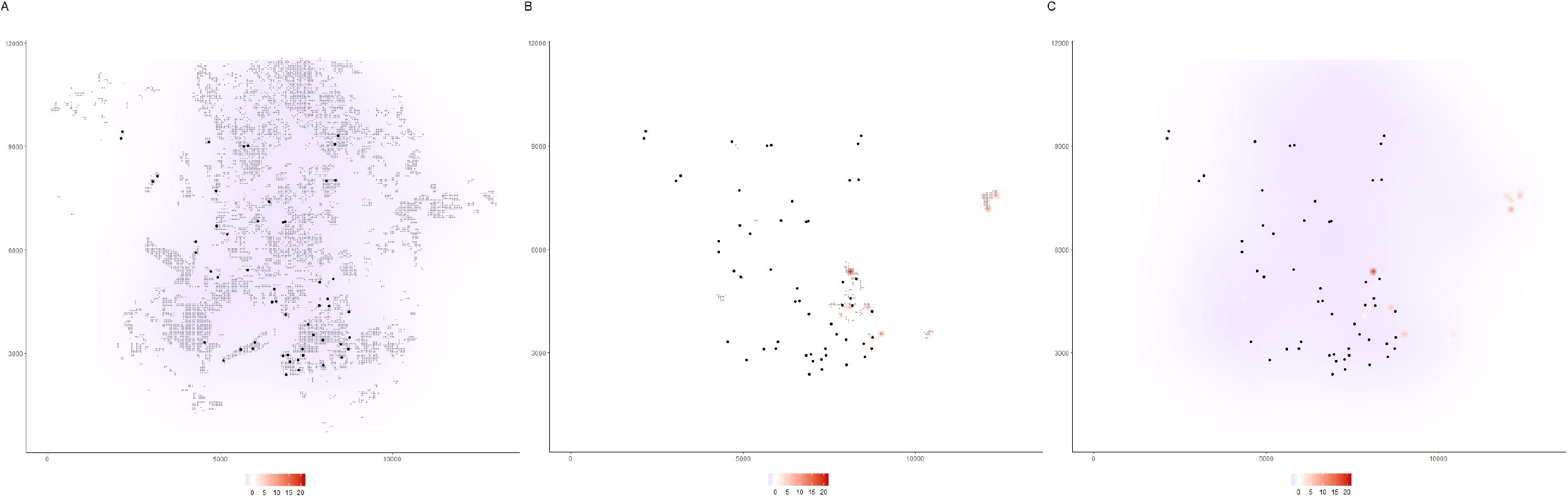
Maps of the predicted effect estimated by the Fsiland method obtained with the function plotFsiland.land() The SIF model was estimated using the command: `resF = Fsiland (Cmoth ~ trait + conv+org, land=landCmoth, data=dataCmoth)`. The three maps A, B and C were obtained with the command `plotFsiland.land (resF, land=landCmoth, data=dataCmoth, var=var), var` equal to `1,2` and `0` respectively. The black points represent locations of response variable observations. In graphics A and B, the gray points are the pixels where conventional and organic orchards are present, respectively. The response surface represents the cumulative effect of the conventional orchards in graphic A, the organic orchards in graphic B and the global landscape in graphic C. At the bottom margin, the bar color gives the effect intensity. In the figure A, conventional orchards had negative effects at large scale. In figure B, organic orchards had positive effects positive effects at small scale. In figure C, global landscape had an overall negative effect except on spotty areas.

## Interpreting spatial influence functions (SIF)

The SIF function describes how the influence of a pixel/cell of a landscape variable is spatially distributed. We assume that the influence is maximal at the pixel location and decreases with the distance. The greater the estimated mean distance of the SIF, the greater the scale of effect of the landscape variable. The estimated SIF can be displayed using the function `plotFsiland.sif()` (see Figure 1). The function `quantile.sif()` allows to quantify the area of medium influence and significant influence of a landscape variable, that we defined as the disc containing 50% and 95% of the influence of the SIF (neglecting 50% and 5% of its broader effect) respectively. They can be compared to the landscape variables distribution in the study area using function `plotFsiland()` (see Figure 1).

## Conclusion

Estimating scales of effect of landscape variables currently remains a great challenge, and consequently so does the estimation of landscape effect. So far no common methodology has emerged. With the siland package, we propose a tool that we believe is useful for all landscape ecologists who wish to investigate this type of question. Here, we have illustrated how with only few siland functions and limited knowledge of R, it is possible to conduct a reproducible and detailed study of multi-scale landscape effect including estimations but also tests and graphs illustrating the results obtained. The “siland” package is very adaptable, it integrates two methods and can handle various types of data. Applications of siland in a multiyear or multisite framework is presented for the buffers method in the `vignette(siland)`. However future developments are still needed to handle the increasing complexity of questions and data (numerous sites, years and dynamic data).

## Authors’ contributions

F.C. and O.M. conceived the ideas and designed methodology, programmed the R package. All authors contributed critically and equally to the drafts and gave final approval for publication.

## Acknowledgements

We are grateful to Claire Lavigne and Pierre Franck for their case study, which illustrates the use of the SILand package, and for their useful advice. We are thankful to Etienne Klein for his relevant comments. This work was financially supported by the Metaprogram SMACH (Sustainable Management of Crop Health; http://www.smach.inra.fr/) of the French National Agronomic Institute (INRA).

