## Supplemental Information 2 for "siland: an R package for estimating the spatial influence of landscape"

### Supplementary information
### Landscape analysis using the R package siland : an application to test the effect of orchards landscape on codling moth density

Florence Carpentier & Olivier Martin

28/02/2020

#### Introduction

We present here an illustration of a landscape analysis using the R package siland. We applied siland methods to analyze the effect of local (treatment) and landscape variables (organic and conventional orchards) on codling moth densities. This example was previously described and analyzed in Ricci et al. (2009) [1]. The first part presents analysis conducted with the Bsiland method (buffers method). The second part presents analysis conducted with the Fsiland method (based on Spatial Influence Function). This analysis was conducted using R version 3.6.2 and package siland version 2.0.

#### Data load

The data are available in the package siland.

```
library(siland)
```

```
## Loading required package: sf
```

```
## Linking to GEOS 3.6.1, GDAL 2.2.3, PROJ 4.9.3
```

```
data(dataCmoth)
data(landCmoth)
```

#### Bsiland method

```
resB=Bsiland(Cmoth~trait+conv+org,land=landCmoth,data=dataCmoth,family="gaussian")
```

```
## Local variables: trait
## Landscape variables: conv org
## Model: Cmoth ~ trait + conv + org
## Model0: Cmoth ~ trait
```

```
#same result with the command Bsiland(Cmoth~trait+conv+org,land=landCmoth,data=dataCmoth)
#argument family is "gaussian" by default
```

[1] Ricci, B., Franck, P., Toubon, J. F., Bouvier, J. C., Sauphanor, B., & Lavigne, C. (2009). The influence of landscape on insect pest dynamics: a case study in southeastern France. Landscape ecology, 24(3), 337-349.

Parameter estimation is based on likelihood maximization. It is therefore strongly recommended to check whether the numerical maximization procedure has converged to the minimum minus log-likelihood value. The Bsiland.lik' function allows to detect whether the estimated minimum is a local mimimun (and not the general minimum).

```
Bsiland.lik(resB,land=landCmoth,data=dataCmoth)
```

```
## Likelihood computing for  conv
## Likelihood computing for  org
```

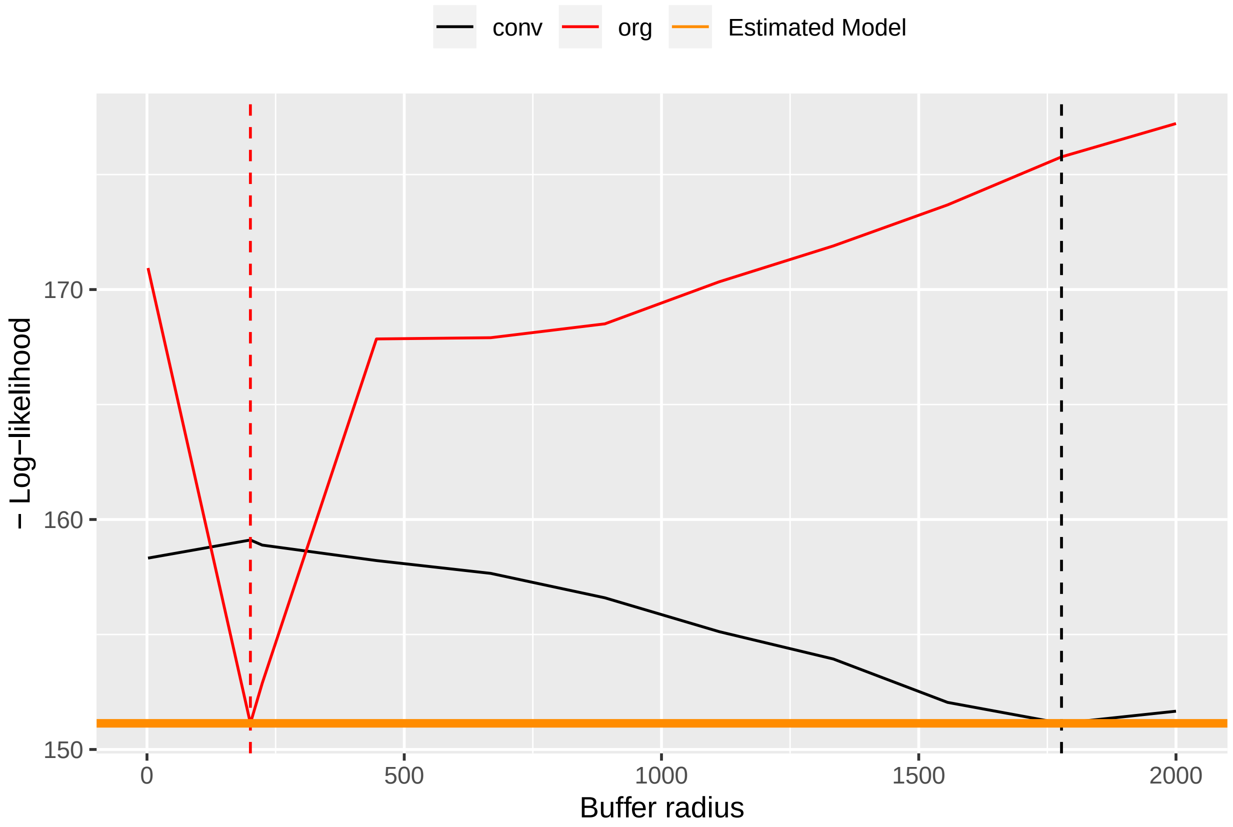


The red and black curves represent the minus log-likelihood in function of the buffers radius of the landscape variables, i.e. the conventional and organic orchards, respectively. The horizontal orange line represents the minus log-likelihood value for estimated parameters. If the estimation proceeded correctly, the red and black curves are minimum when they reach the orange horizontal line (see `vignette(siland)` for more details). This was the case here.

By printing Bsiland's result object, we obtained the estimated parameters and the global test of landscape effects (i.e. H0: "No landscape variable has an effect" vs H1: "At least one of the landscape variables has an effect").

```
resB
```

```
## Model: Cmoth ~ trait + conv + org
##
## Landscape variables: conv org
##
## Coefficients:
## (Intercept)       trait        conv         org      B.conv       B.org
##      11.222      -0.155     -40.913      87.566    1777.529     201.021
```

```
##
## standard error: 4.130343
## AIC: 316.27  AIC (no landscape): 364.79
## (No landscape effect) p-value: 1.562683e-11
```

Here the landscape has a global significant effect (p.val= 1.562683e-11).

The function `summary()` provides significance tests of the intensity of the effects of the explanatory variables (local or landscape).

```r
summary(resB)
```

```
## Buffer sizes:
##    B.conv     B.org
## 1777.5294  201.0211
##
## -- Tests are given conditionnaly to the best estimated buffer sizes --
##
## Call:
## Cmoth ~ trait + conv + org
##
## Deviance Residuals:
##      Min        1Q    Median        3Q       Max
## -13.1294   -1.4184   -0.1032    1.8552   14.8693
##
## Coefficients:
##             Estimate Std. Error t value Pr(>|t|)
## (Intercept)  11.2225     2.7566   4.071 0.000167 ***
## trait        -0.1550     0.1507  -1.028 0.308725
## conv        -40.9131     9.2650  -4.416 5.39e-05 ***
## org          87.5657     9.2722   9.444 1.06e-12 ***
## ---
## Signif. codes:  0 '***' 0.001 '**' 0.01 '*' 0.05 '.' 0.1 ' ' 1
##
## (Dispersion parameter for gaussian family taken to be 17.05973)
##
##     Null deviance: 2867.13  on 53  degrees of freedom
## Residual deviance:  852.99  on 50  degrees of freedom
## AIC: 312.27
##
## Number of Fisher Scoring iterations: 2
```

The buffer sizes for conventional and organic orchards were estimated at 1777.52 m and 201.02 m, respectively. The effect of the local variable was estimated at -0.1550 but not significant (p.val=0.308734). The intensity of the effect of conventional orchards in buffers of size 1777.52 m was estimated negative (-40.9131) and significant (p.val<0.0001). The intensity of the effect of organic orchards in buffers of size 201.02 m was estimated positive (87.5657) and significant (p.val<0.0001).

Optimal radii for landscape variables are stored in 'resB$parambuffer':

```r
resB$parambuffer
```

```
##    B.conv     B.org
## 1777.5294  201.0211
```

and percentages of each lansdcape variable on each associated observations buffers are stored in 'resB$buffer':

```r
head(resB$buffer)
```

```
##        conv        org
## 1 0.2564024 0.04015922
## 2 0.2654119 0.00000000
## 3 0.1982562 0.04295240
## 4 0.1891139 0.00000000
## 5 0.2651649 0.00000000
## 6 0.2622895 0.00000000
```

A graphical representation of the estimated buffers can be obtained with the function 'plotBsiland.land'. The argument 'var' indicates the indice of the plotted landscape variable.

The map of the negative effect of conventional orchards was obtained by the following line code :

```r
plotBsiland.land(resB,land=landCmoth,data=dataCmoth,var=1)
```

```
## Plot for landscape variable  conv
##   B.conv
## 1777.529
```

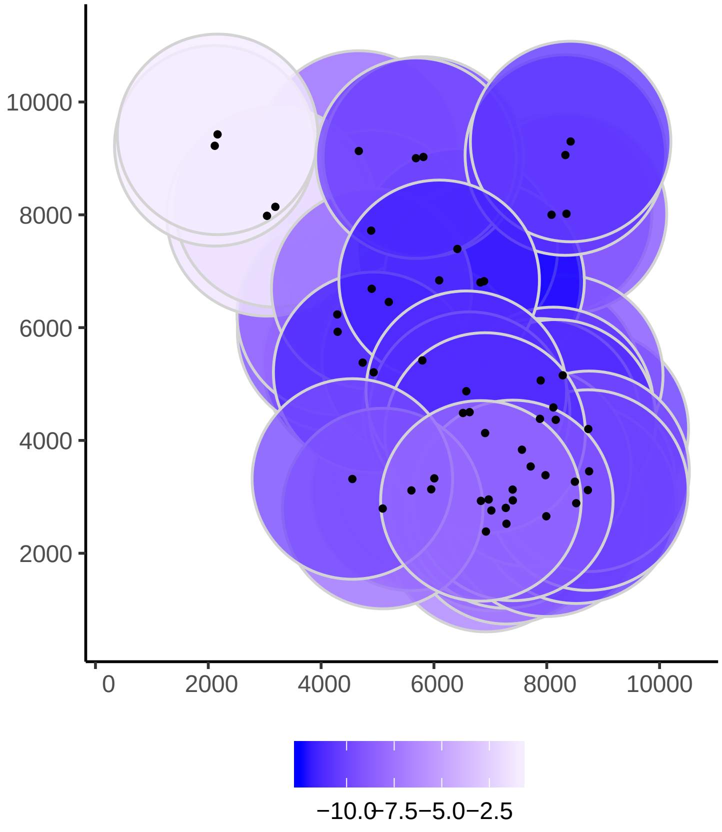


The map of the positive effect of organic orchards was obtained by the following line code :

```r
plotBsiland.land(resB,land=landCmoth,data=dataCmoth,var=2)
```

```
## Plot for landscape variable  org
##    B.org
## 201.0211
```

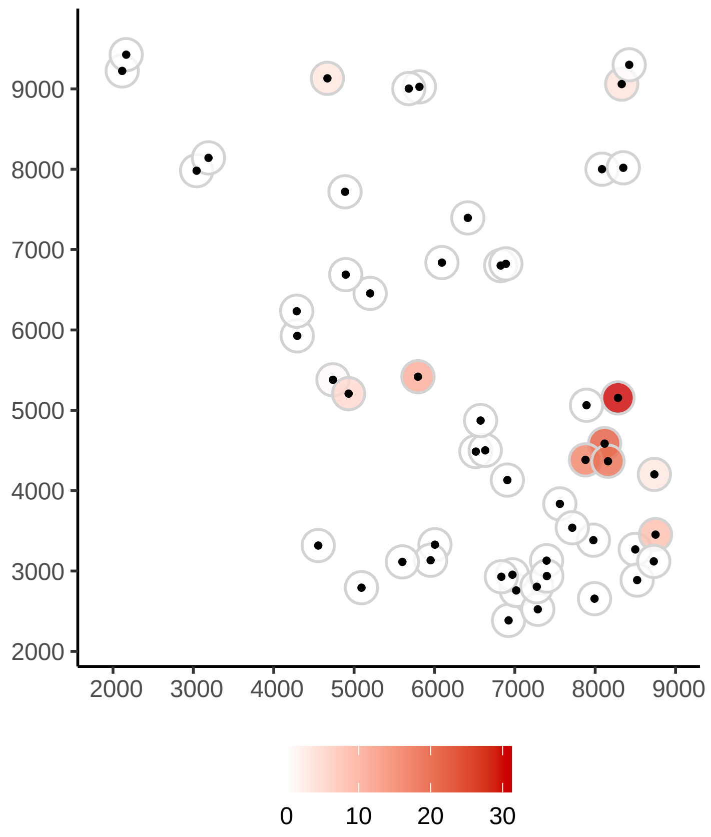


The following commands allow to compute AIC, BIC, predicted values and residuals of the model :

```
AIC(resB)
```

```
## AIC =  316.272
```

```
BIC(resB)
```

```
## BIC =  330.195
```

```
head(fitted(resB))
```

```
##         1         2         3         4         5         6
##  2.699215  0.363623  5.632660  2.245547 -1.175857 -1.058215
```

```
head(residuals(resB))
```

```
##         1         2         3         4         5         6
## -2.439956 -0.363623 -4.465993 -2.145547  1.604428  1.858215
```

We can consider that the effect of conventional and organic orchards did not start from the observation sites, but from the boundary of the orchard where the observations are located using the “border=T” argument.

```
resB2=Bsiland(Cmoth~trait+conv+org,land=landCmoth,data=dataCmoth,border=T)
```

```
## Local variables: trait
## Landscape variables: conv org
## Model: Cmoth ~ trait + conv + org
## Model0: Cmoth ~ trait
```

```r
resB2
```

```
## Model: Cmoth ~ trait + conv + org
##
## Landscape variables: conv org
##
## Coefficients:
## (Intercept)       trait        conv         org      B.conv       B.org
##       2.418       0.098      -6.186      72.182     273.900     126.984
##
## standard error: 4.96772
## AIC: 336.21  AIC (no landscape): 364.79
## (No landscape effect) p-value: 2.198806e-07
```

```r
summary(resB2)
```

```
## Buffer sizes:
##   B.conv    B.org
## 273.9003 126.9844
##
## -- Tests are given conditionnaly to the best estimated buffer sizes --
##
## Call:
## Cmoth ~ trait + conv + org
##
## Deviance Residuals:
##      Min        1Q    Median        3Q       Max
## -18.8039   -1.3303   -0.3306    1.1690   16.7261
##
## Coefficients:
##             Estimate Std. Error t value Pr(>|t|)
## (Intercept)  2.41823    2.16282   1.118    0.269
## trait        0.09811    0.17047   0.576    0.568
## conv        -6.18557    3.91884  -1.578    0.121
## org         72.18227   11.75055   6.143 1.31e-07 ***
## ---
## Signif. codes:  0 '***' 0.001 '**' 0.01 '*' 0.05 '.' 0.1 ' ' 1
##
## (Dispersion parameter for gaussian family taken to be 24.67824)
##
##     Null deviance: 2867.1  on 53  degrees of freedom
## Residual deviance: 1233.9  on 50  degrees of freedom
## AIC: 332.21
##
## Number of Fisher Scoring iterations: 2
```

```r
plotBsiland.land(resB2,land=landCmoth,data=dataCmoth,var=1)
```

```
## Plot for landscape variable  conv
##   B.conv
## 273.9003
```

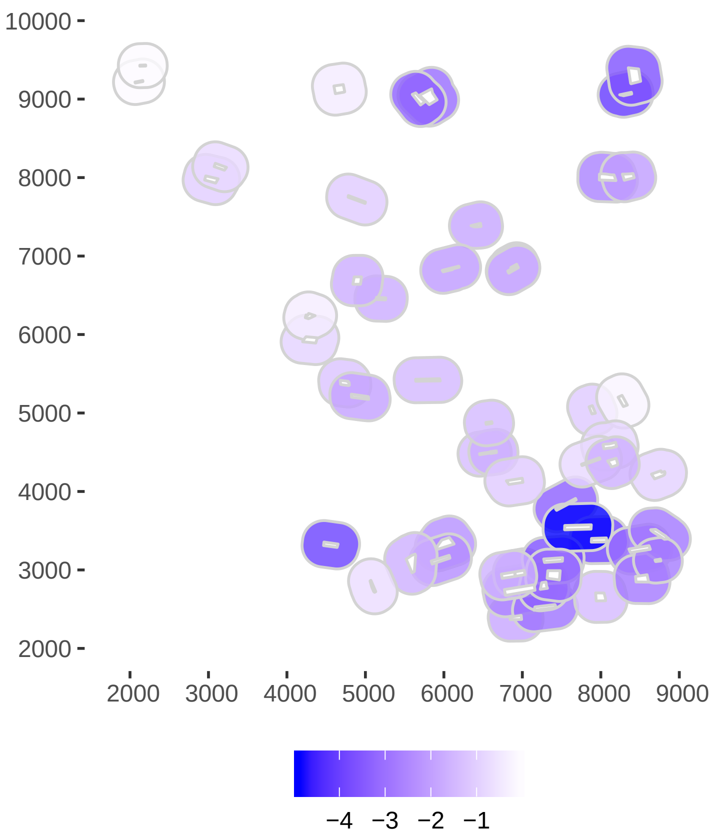


```r
plotBsiland.land(resB2,land=landCmoth,data=dataCmoth,var=2)
```

```
## Plot for landscape variable  org
##    B.org
## 126.9844
```

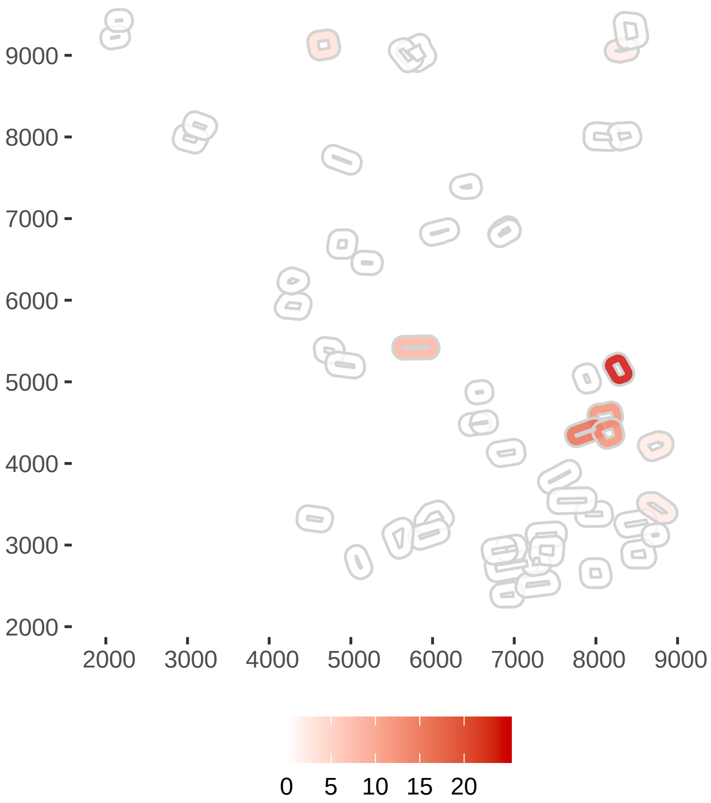


#### Fsiland method

In the approach Fsiland, Spatial Influence Functions (SIF) are used to modelize the influence of landscape variables (decreasing with distance). In this approach, the scale of effect of a landscape variable is estimated by the parameter of the SIF.

```
resF=Fsiland(Cmoth~trait+conv+org,land=landCmoth,data=dataCmoth)
```

```
## Local variables: trait
## Landscape variables: conv org
## Model: Cmoth ~ trait + conv + org
## Model0: Cmoth ~ trait
```

```
# equivalent to resF=Fsiland(Cmoth~trait+conv+org,land=landCmoth,data=dataCmoth,family="gaussian")
```

Using 'Fsiland.lik' we checked the convergence of the estimation procedure :

```
Fsiland.lik(resF,land=landCmoth,data=dataCmoth)
```

```
## Likelihood computing for  conv
## Likelihood computing for  org
```

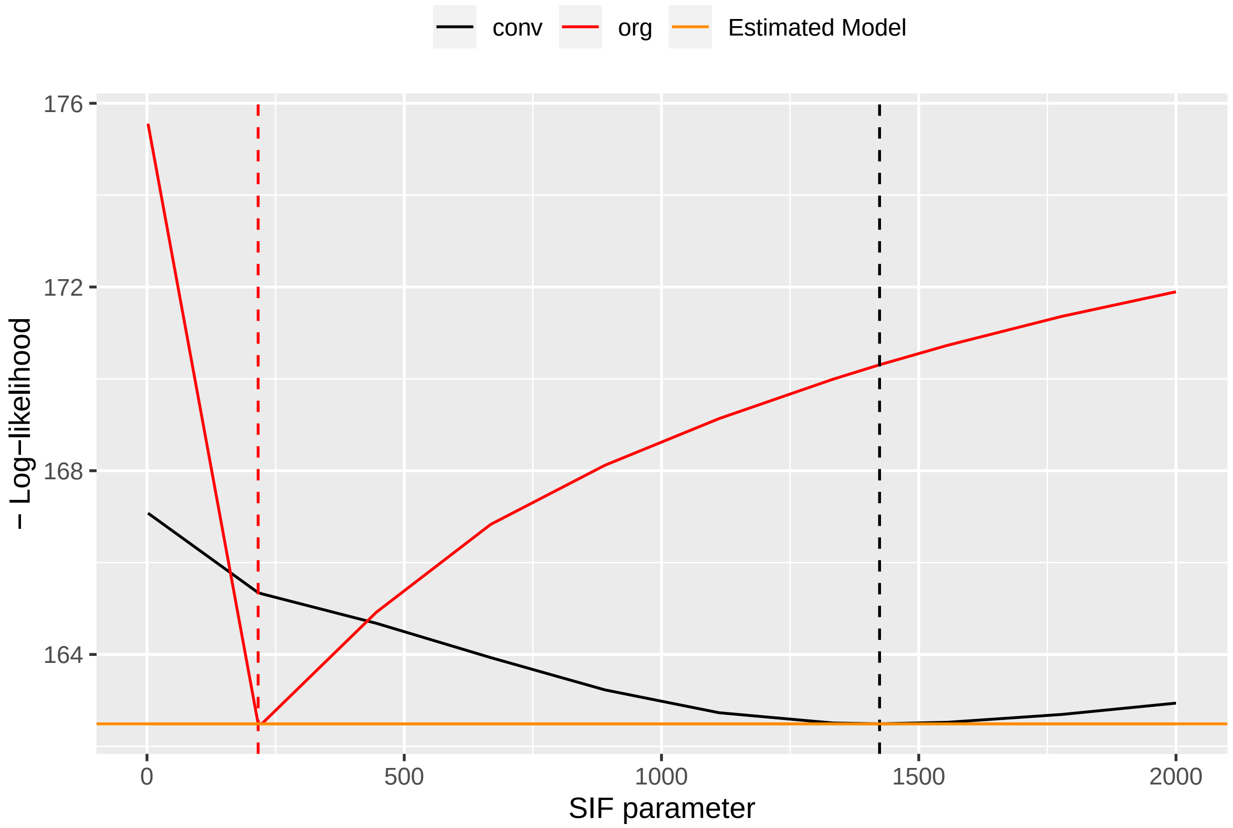


As the red and black curves were minimum when they reached the orange horizontal line, the estimation seemed to proceed correctly.

By printing the Fsiland's result object, we obtained the estimated parameters and the global test of landscape effects (i.e. H0: "No landscape variable has an effect" vs H1: "At least one of the landscape variables has an effect").

```
resF
```

```
## Model: Cmoth ~ trait + conv + org
##
## Landscape variables: conv org
##
## Coefficients:
## (Intercept)       trait        conv         org    SIF.conv     SIF.org
##       9.201      -0.013     -11.856      24.880    1423.903     216.085
##
## standard error: 5.096762
## AIC: 338.98  AIC (no landscape): 364.79
## (No landscape effect) p-value: 8.15168e-07
```

The function `summary()` provides parameters estimation and significance tests of the intensity of the effects of explanatory variables (local or landscape).

```
summary(resF)
```

```
## SIF parameters:
##  SIF.conv   SIF.org
## 1423.9033  216.0852
```

```
##
## -- Tests are given conditionnaly to the best SIF parameters --
##
## Call:
## Cmoth ~ trait + conv + org
##
## Deviance Residuals:
##      Min        1Q    Median        3Q       Max
## -17.0653   -1.6312   -0.0058    1.0128   24.1851
##
## Coefficients:
##             Estimate Std. Error t value Pr(>|t|)
## (Intercept)   9.2008     3.3927   2.712  0.00915 **
## trait        -0.0129     0.1842  -0.070  0.94446
## conv        -11.8561     3.8669  -3.066  0.00349 **
## org          24.8800     3.9593   6.284 7.91e-08 ***
## ---
## Signif. codes:  0 '***' 0.001 '**' 0.01 '*' 0.05 '.' 0.1 ' ' 1
##
## (Dispersion parameter for gaussian family taken to be 25.97699)
##
##     Null deviance: 2867.1  on 53  degrees of freedom
## Residual deviance: 1298.8  on 50  degrees of freedom
## AIC: 334.98
##
## Number of Fisher Scoring iterations: 2
```

The mean distances for SIF of conventional and organic orchards were estimated at 1423.9033 m and 216.0852 m, respectively. The effect of the local variable treatment was estimated at-0.012 but not significant (p.val=0.94446). The intensity of the effect of conventional orchards was estimated negative (-11.8561) and significant (p.val=0.00349). The intensity of the effect of organic orchards was estimated positive (24.8800) and significant (p.val<0.001).

The map of the effects of landscape variable can be obtain using `plotFsiland.land`. The argument 'var' indicates the indice of the considered landscape variable. For plotting effects of conventional orchard :

```
plotFsiland.land(resF,land=landCmoth, data=dataCmoth,var=1)
```

```
## [1] "Distance computing... Wait..."
## [1] "Contribution computing... Wait.."
```

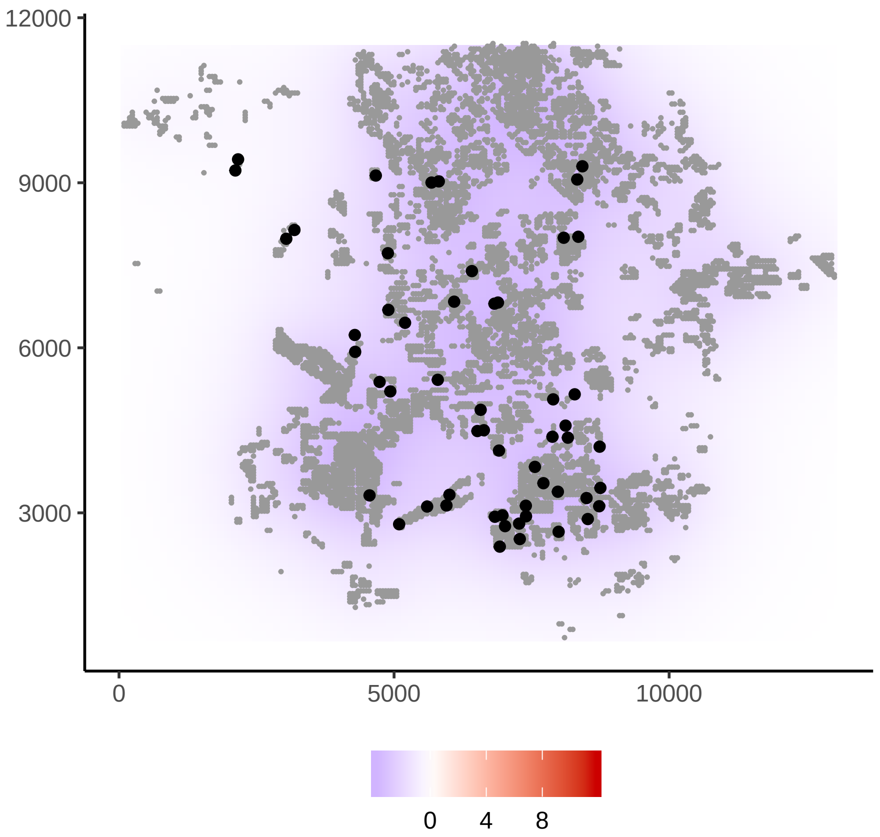


For plotting effects of organic orchards :

```r
plotFsiland.land(resF,land=landCmoth, data=dataCmoth,var=2)
```

```
## [1] "Distance computing... Wait..."
## [1] "Contribution computing... Wait.."
```

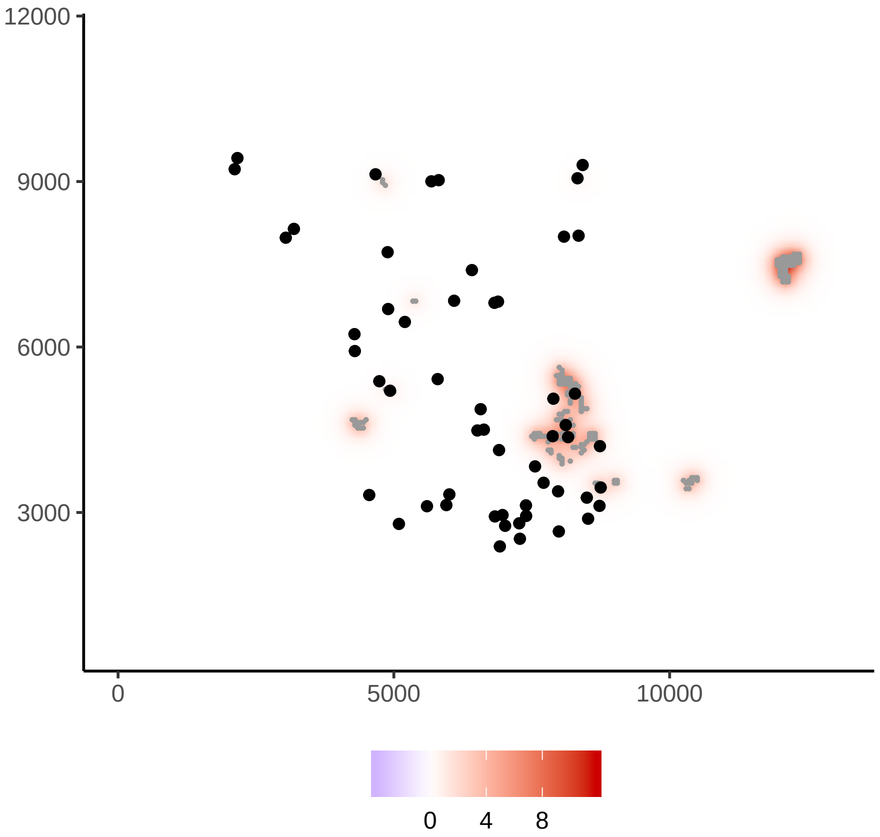


The map of the global landscape effect i.e. the sum of the effects of all landscape variables is obtained using `var=0`,

```
plotFsiland.land(resF,land=landCmoth, data=dataCmoth,var=0)
```

```
## [1] "Distance computing... Wait..."
## [1] "Contribution computing... Wait.."
```

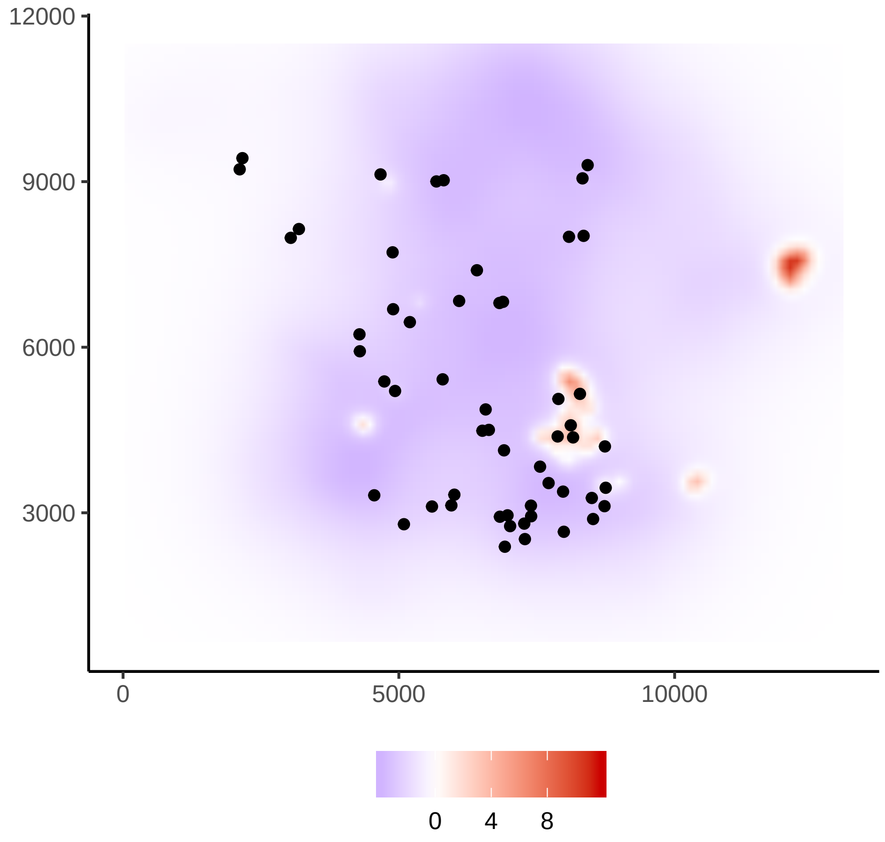


The estimated SIF can be plotted using the function `plotFsiland.sif`:

```
plotFsiland.sif(resF)
```

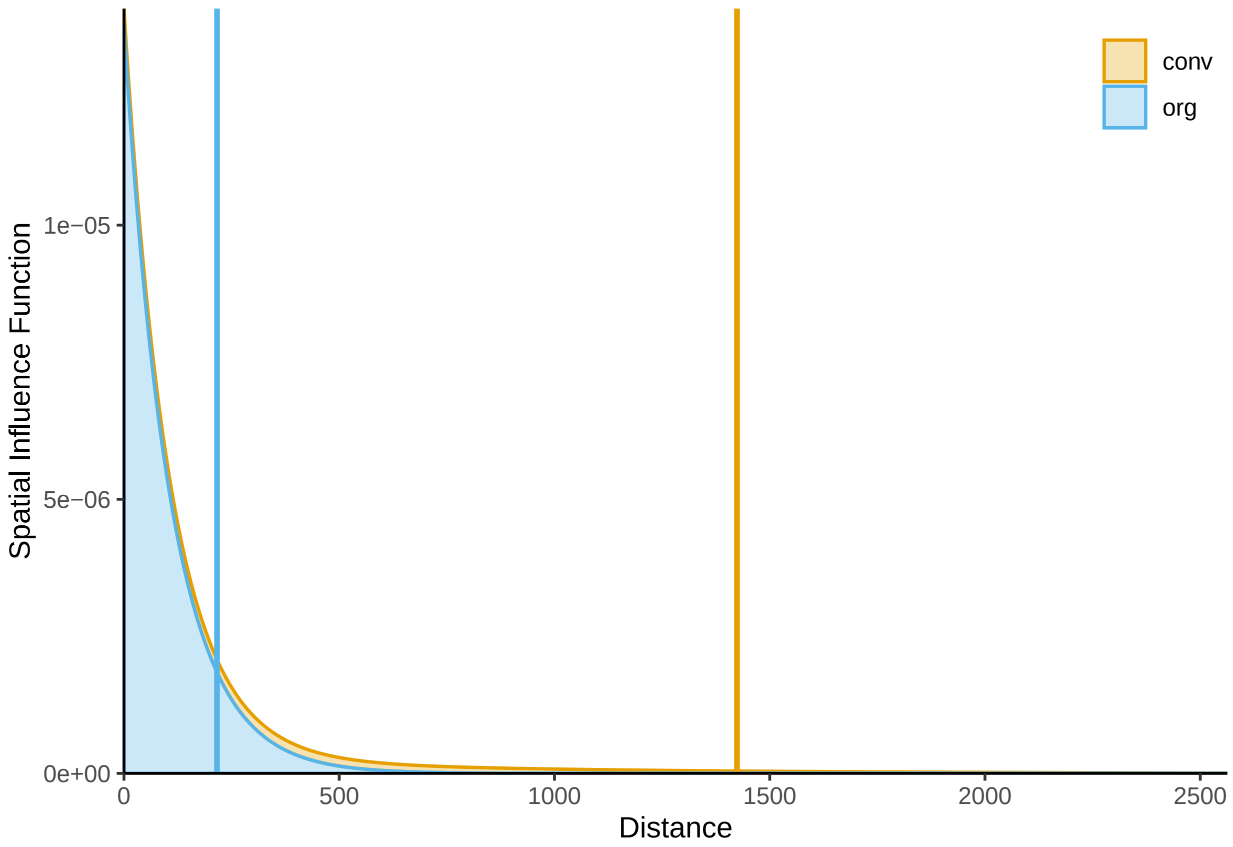


The observations map and the significant and medium effect area associated to the estimated SIFs can be plotted using the function `plotFsiland.sif`:

```
plotFsiland(resF,landCmoth, data=dataCmoth)
```

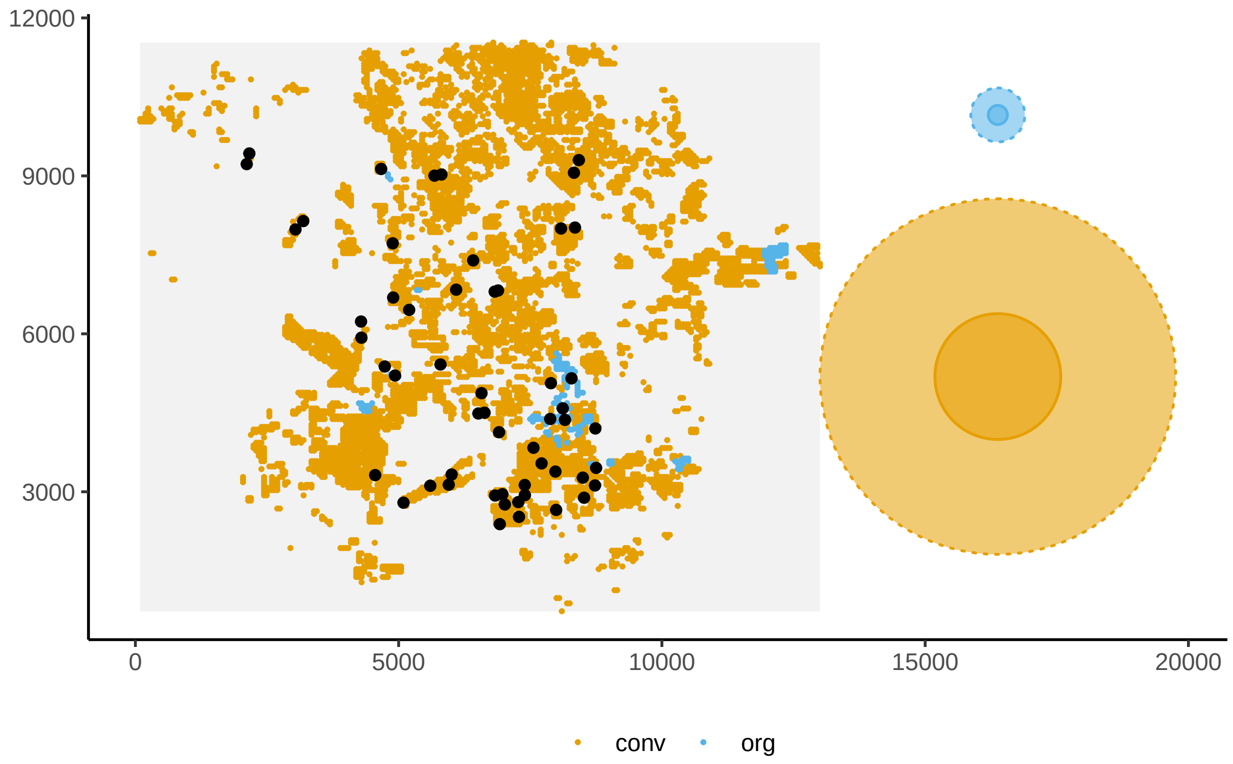


We can consider that the effect of conventional and organic orchards do not start from the observation locations but from the border of the orchard where observations are located using the argument 'border=T'.

```
resF2=Fsiland(Cmoth~trait+conv+org,land=landCmoth,data=dataCmoth,family="gaussian",border=T)
```

```
## Local variables: trait
## Landscape variables: conv org
## Model: Cmoth ~ trait + conv + org
## Model0: Cmoth ~ trait
```

```
summary(resF2)
```

```
## SIF parameters:
##  SIF.conv   SIF.org
## 1496.5480  200.8924
##
## -- Tests are given conditionnaly to the best SIF parameters --
##
## Call:
## Cmoth ~ trait + conv + org
##
## Deviance Residuals:
##      Min        1Q    Median        3Q       Max
## -19.7059   -1.5027    0.0229    1.2009   19.1245
##
## Coefficients:
##                Estimate Std. Error t value Pr(>|t|)
## (Intercept)     8.25920    3.27067   2.525  0.01478 *
```

```
## trait          0.01061    0.17504   0.061  0.95191
## conv         -11.52326    3.97911  -2.896  0.00559 **
## org           38.05239    5.55268   6.853 1.02e-08 ***
## ---
## Signif. codes:  0 '***' 0.001 '**' 0.01 '*' 0.05 '.' 0.1 ' ' 1
##
## (Dispersion parameter for gaussian family taken to be 24.35624)
##
##     Null deviance: 2867.1  on 53  degrees of freedom
## Residual deviance: 1217.8  on 50  degrees of freedom
## AIC: 331.5
##
## Number of Fisher Scoring iterations: 2
```

```
plotFsiland.land(resF2,land=landCmoth, data=dataCmoth,var=0)
```

```
## [1] "Distance computing... Wait..."
## [1] "Contribution computing... Wait.."
```

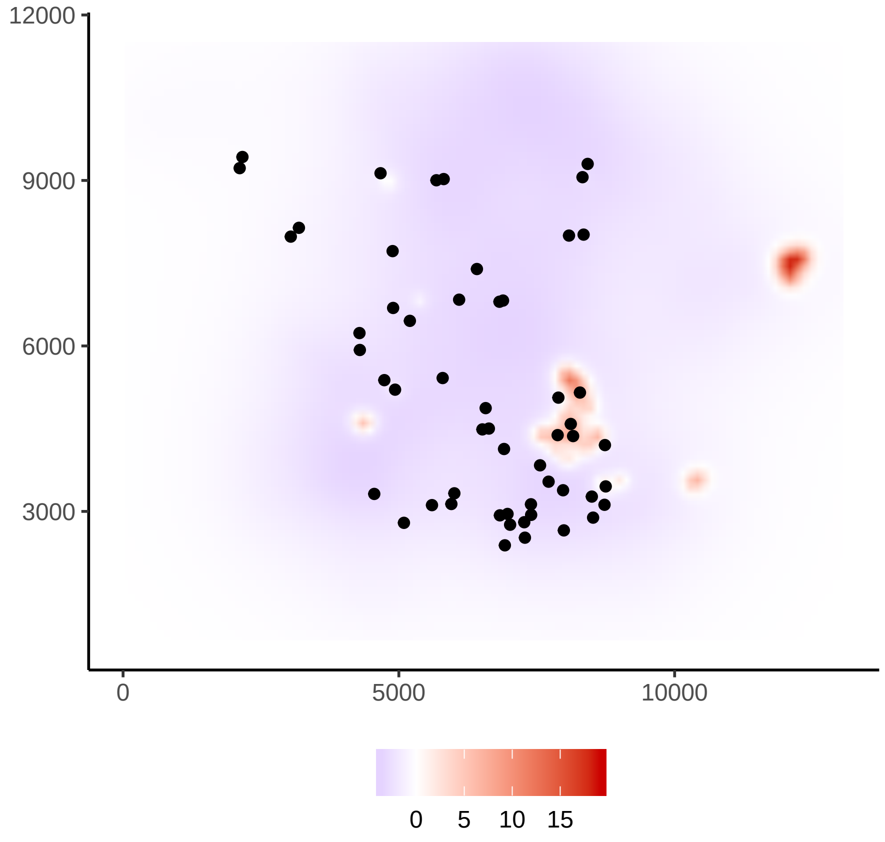
